# Membrane Interactions of α-Synuclein Revealed by Multiscale Molecular Dynamics Simulations, Markov State Models, and NMR

**DOI:** 10.1101/2020.06.18.156216

**Authors:** Sarah-Beth T. A. Amos, Thomas C. Schwarz, Jiye Shi, Benjamin P. Cossins, Terry S. Baker, Richard J. Taylor, Robert Konrat, Mark S. P. Sansom

## Abstract

α-Synuclein is a presynaptic protein that binds to cell membranes and is linked to Parkinson’s disease (PD). Whilst the normal function of remains α-synuclein remains uncertain, it is thought that oligomerization of the protein on the cell membrane contributes to cell damage. Knowledge of how α-synuclein binds to lipid bilayers is therefore of great interest as a likely first step in the molecular pathophysiology of PD, and may provide insight of the phenotype of PD-promoting mutations. We use coarse-grained and atomistic simulations in conjunction with NMR and cross-linking mass spectrometry studies of α-synuclein bound to anionic lipid bilayers to reveal a break in the helical structure of the NAC region, which may give rise to subsequent oligomer formation. Coarse-grained simulations of α-synuclein show that the interhelical region leads recognition and binding to both POPG and mixed composition bilayers and identifies important protein-lipid contacts, including those in the region between the two helices in the folded structure. We extend these simulations with all-atom simulations of the initial binding event to reveal details of the time-progression of lipid binding. We present secondary structure analysis that reveals points of possible β-strand formation in the structure, and investigate intramolecular contacts with simulations and mass-spectrometry crosslinking. Additionally we show how Markov state models can be used to investigate possible conformational changes of membrane bound α-synuclein in the NAC region, and we extract representative structures. These structural insights will aid the design and development of novel therapeutic approaches.

## Introduction

α-Synuclein is a protein implicated in neurodegenerative disorders including Parkinson’s disease and Lewy body dementia (1). Its function in healthy neurons remains uncertain (2). Lipid bilayer association of α-synuclein is thought to be important for its biological function in regulating synaptic vesicles, where it has been shown to be essential for SNARE complex assembly at the presynaptic membrane (3, 4). It has been postulated to have a wide range of functions, including neuronal differentiation (4) and suppression of apoptosis (5). It is thought that toxicity towards neurones arises from the interaction of misfolded/aggregated α-synuclein with the lipid bilayer component of cell membranes (6). Thus, defining the interactions between α-synuclein and lipid bilayers is a key step in understanding the mode of action of α-synuclein, thus helping to provide a route towards drug research aimed at presenting or reversing its cellular effects.

Native α-synuclein is an intrinsically disordered monomeric protein in solution (7-9). SAXS and NMR-derived experimental data suggest relatively small amounts of compaction in the ensemble of free α-synuclein (10, 11). In contrast to these observations some discrete molecular dynamics (MD) simulations (12) combined with crosslinking data (13) have been interpreted as suggesting that part of the ensemble may be relatively compact. Partial secondary structure in the free state of α-synuclein is observed experimentally in solid-state NMR measurements (14) and in the computational discrete MD approach. (12) Solution and solid state NMR, EPR FRET and CD studies reveal that α-synuclein folds into a predominantly α-helical structure when it interacts with SDS micelles or liposomes, the surface of which provide a mimic of negatively charged lipid bilayer membranes (14-19). The NMR structure when bound to SDS micelles (PDB ID 1XQ8; Fig. 1A) (17, 20) is formed by two α-helices (residues 1-38 and residues 46-95) in the N-terminal section of the molecule followed by a disordered C-terminal segment (residues 98 to 140). The α-helical N-terminal segment of α-synuclein contains seven KTKEGV motifs which are thought to play a role in maintaining the native unfolded status in solution (21). The disordered C-terminal tail contains 10 glutamate and 5 aspartate residues, and so is highly negatively charged. This distribution of charged residues helps to explain the observed pattern of secondary structure formation upon binding to an anionic micelle.

**Figure 1:**
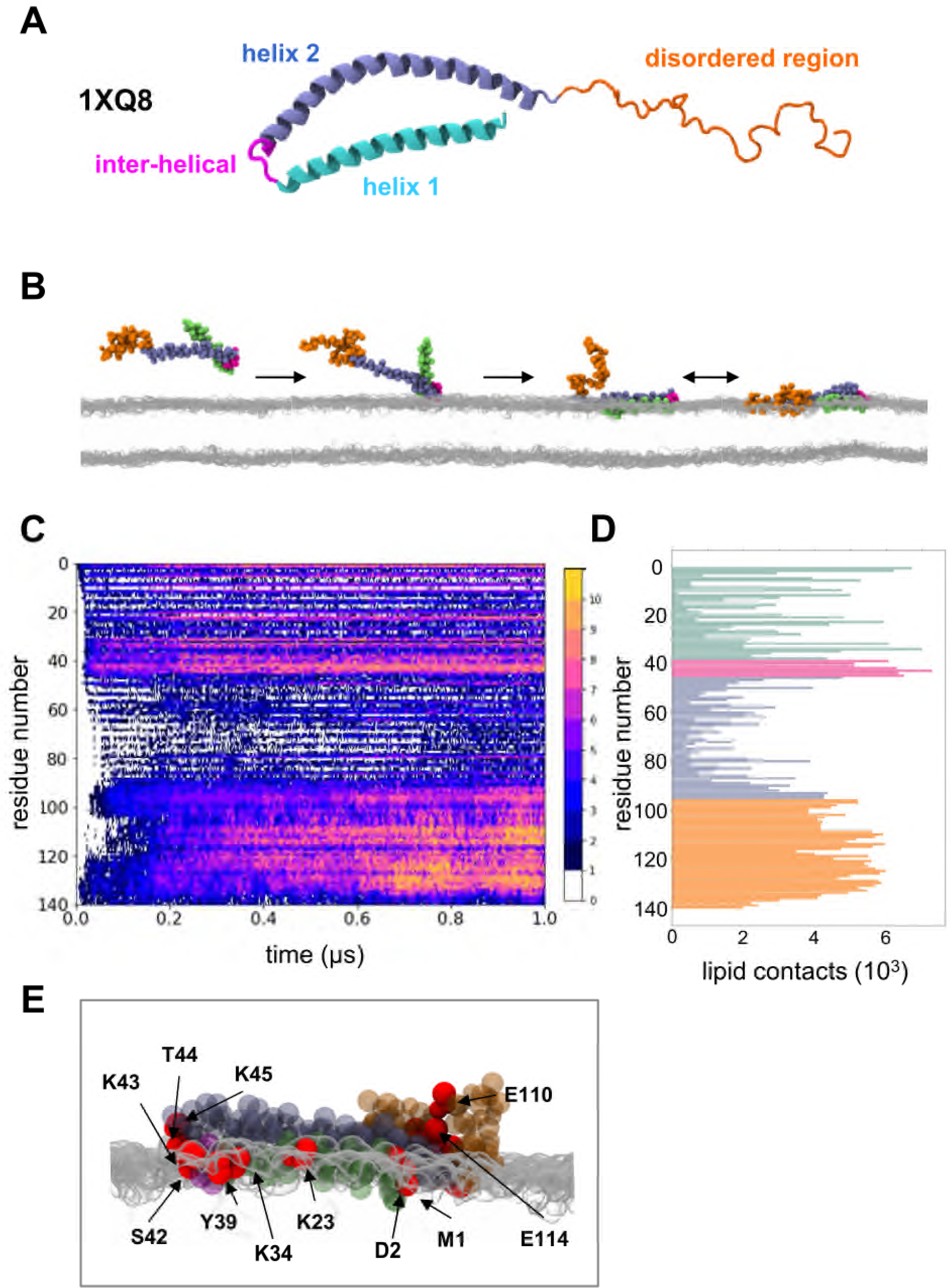
The structure of α-synuclein and its interaction with an anionic lipid bilayer. **A** The folded α-synuclein structure (PDB id 1XQ8) contains three regions: two helices (1 cyan and 2 grey/purple) separated by an interhelical loop (pink) and followed by a C-terminal disordered region (orange). **B** Successive snapshots from a CG simulation of the interaction of α-synuclein with a PG bilayer. Monomeric α-synuclein is initially positioned distal to the membrane. During 10 replicate simulations each of 1 μs duration α-synuclein bound to the membrane. Colours as in **A. C** Total number of simulations across the ensemble where each residue of α-synuclein is in contact with the membrane. **D** Total number of individual contacts of α-synuclein with PG summed over the 10 simulation replicates. A residue is considered in contact with the lipid if the residue backbone particle is within 0.7 nm of any lipid particle. **E** Side view of α-synuclein bound to the surface of a PG membrane from the endpoint of a simulation. Red particles indicate the top 14 residues making contact with PG lipid headgroups shown in grey.

The relevance of lipid interactions to the function of α-synuclein is demonstrated by its structural rearrangement upon vesicle binding (15), and by the retention of the overall structure in Parkinson disease variants as seen by NMR (20). Furthermore, disease causing mutations alter lipid-induced generation of fibrils (22). There is some evidence that oligomeric intermediates, which form only on the membrane, are more toxic than the mature fibrils (23-26). In this context, it becomes important to understand the initial binding event of α-synuclein to the lipid membrane, alongside subsequent conformational rearrangements that may take place on the bilayer surface. Furthermore, the modulation of membrane binding as a therapeutic strategy remains an open question (27).

Computational approaches, and in particular MD, have played a key role in exploring the conformational dynamics of α-synuclein and related proteins, both in aqueous solution (28-32) and when bound to membranes (33-39). MD simulations have also been shown to be a powerful tool for characterising the interactions of proteins with membrane and lipids (40, 41). Here we use a multiscale approach, combining coarse-grained and atomistic MD simulations, to explore the initial interaction of α-synuclein with anionic lipids bilayers, and to describe secondary structure stabilization of the membrane-bound protein.

## Results & Discussion

In order to investigate the interactions of α-synuclein with model membranes, we adopted a multiscale approach combining coarse-grained (CG) simulations of the protein/bilayer encounter followed by atomistic (AT) simulations to explore conformational changes of the protein when bound to the bilayer surface. Simulations are summarized in Table 1, and details are provided in the Methods.

**Table 1:** Summary of Simulations.

| description | CG/AT* | lipids | time<br>(μs) | $N$ | total<br>(μs) |
| --- | --- | --- | --- | --- | --- |
| α-Syn/bilayer | CG | POPG | 1 | 10 | 10 |
| α-Syn/bilayer, varying helicity | CG | POPG | 1 | 4x10 | 40 |
| α-Syn/mixed lipid bilayer | CG | DOPC/DOPE/DOPS<br>2:5:3 | 2 | 10 | 20 |
| Conformational changes | AT | POPG | 0.1 to<br>0.25 | 3x10 | 5.5 |

\*CG = coarse grained; AT = atomistic

### CG simulations reveal the initial membrane binding mode of α-synuclein

CG simulations were set up by positioning the α-synuclein monomer (in a CG representation corresponding to the structure in the presence of micelles, i.e. PDB id 1XQ8, Fig. 1AB) 4 nm away from an anionic POPG membrane surface (Fig.1B). The protein molecule diffused towards and bound to the membrane surface within less than 1 μs. These simulations revealed that the interhelical region formed the first contacts with the POPG membrane (Fig.1BC). Thus, 50% of simulations showed binding of the interhelical region within ∼0.1 μs, and in all simulations the protein bound to the bilayer within 1 μs. Residues in the N-terminal Helix 1 (residues 1-38) forming multiple contacts with POPG headgroups included residues 1, 21, 23, 32 and 34. Helix 2 (residue 46-95) formed few contacts with the POPG membrane, although residues 50 and 80 contributed to the overall binding profile as the simulation progresses. The disordered C-terminus makes a number of contacts with the membrane as the simulation progresses (Fig. 1CD) This correlates with experimental observation of calcium-mediated contacts of the C-terminus to synaptic vesicles (42).

In order to further analyse the POPG binding profile, we conducted correlation analysis of the contacts, which shows strong positive correlation in contacts between the disordered tail and the interhelical region and residues 79-80 (SI Fig. S1A). Principal components analysis revealed that 80 residues explain 95% of the variance in binding (SI Fig. S1B). A representative bound structure of α-synuclein with the top 14 residues making lipid contacts highlighted in red is shown in Fig. 1E (and in SI Fig. S1C). Four of these residues (39, 42, 43, 44, 45) are in the interhelical region. Overall, we observed that Helix 1 formed close contacts with the membrane, with major contributions to the contact binding profile from residues 1, 2, and 32. Interestingly, we observe that Helix 2 (containing the NAC region) is folded back over the top of Helix 1.

In the CG protocol, the secondary structure present in the initial structure (PDB id 1XQ8) is maintained by restraints during the simulation. To establish how variation in the secondary structure of α-synuclein might contribute to the binding profile, we modelled the α-synuclein monomer with decreasing degrees of secondary structure restraints: residues 1-80, 1-60, 1-40, and 1-20 (Table 1 and SI Fig. S2). As the restraints were relaxed, we observed a greater contribution to the contact profile from Helix 2. The large contribution from the interhelical region remains unchanged. The increased binding contribution from the Helix 2 region reflects its greater conformational freedom to make contacts with the bilayer instead of being folded over Helix-1 (as in Fig.1C). The interhelical region appears to dominate the contact profile even when the restraints are reduced.

We also explored the binding of the α-synuclein monomer to a more complex model membrane. The mixed lipid bilayer employed (DOPE/DOPS/ DOPC in a 5:3:2 molar ratio) was intended to provide a simple mimic of a synaptic vesicle membrane. Thus is still anionic but with 30% of the net surface charged of the POPG bilayer used previously. As can be seen from Fig.2A the initial interaction remains led by the interhelical region, with a reduced contribution from the unstructured C-terminus. Analysing the contribution of each lipid species to the contact profile (Fig. 2BC) shows that interactions with DOPE and DOPS are preferred to those with DOPC, as has been observed for a number of experimental studies (43). Overall these simulations confirm those with POPG in indicating the importance of Helix-1 and the interhelical region. A number of known natural mutations are found in these two regions, including A30P and E46K both of which are associated with differentiated disease pathology (22).

**Figure 2:**
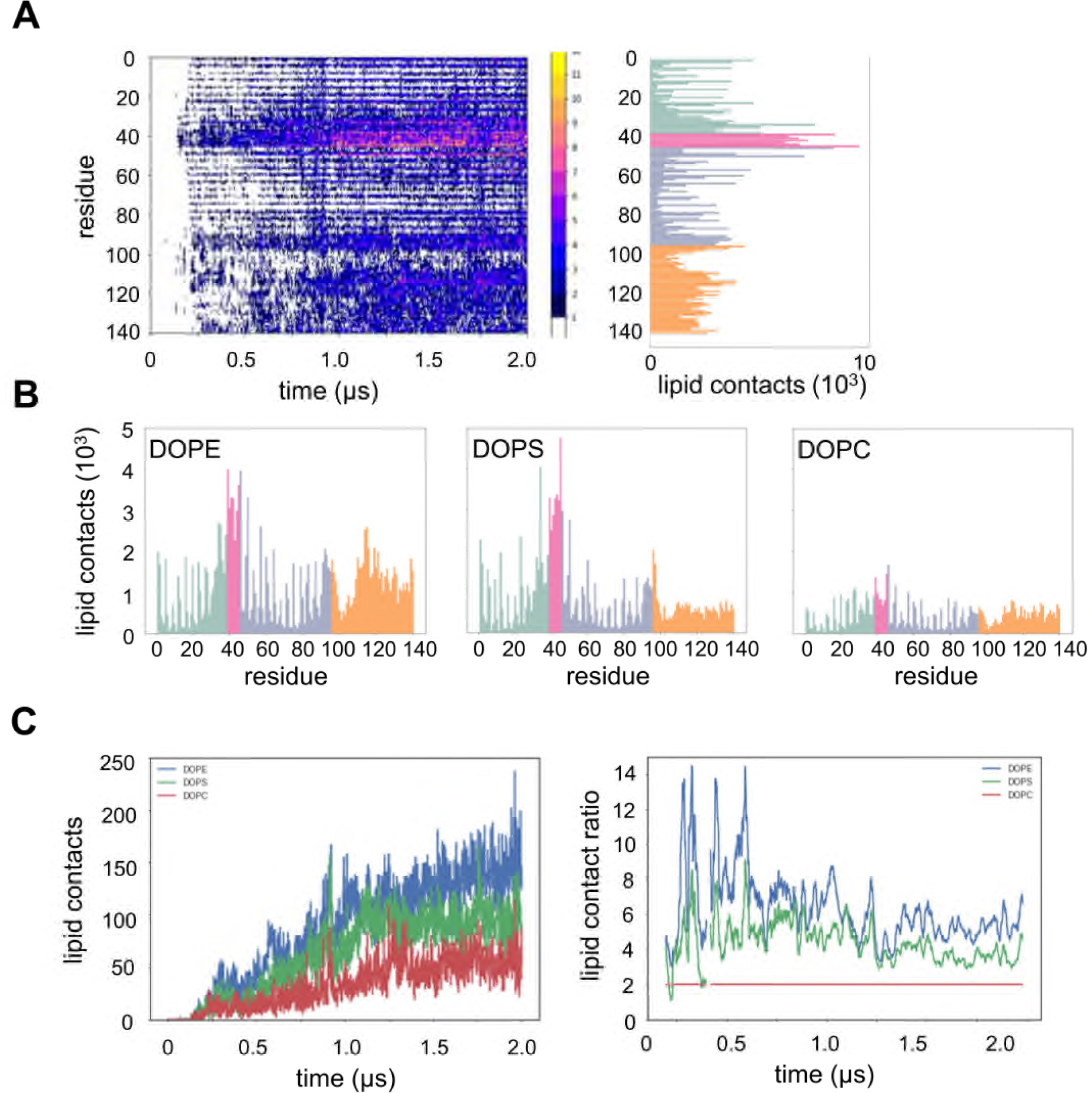
Interaction of α-synuclein with a PC/PE/PS (2:5:3) lipid bilayer. Simulations were of 10 replicates each for 2 *μ*s. **A** Contacts for with all lipids for each residue of α-synuclein summed over the 10 simulation replicates. Colours on the histogram indicate the structural regions defined in Fig. 1. **B** Total contacts shown separately for the three lipid species. **C** Total contacts for each lipid species (PC = red; PE = blue; PS = green) shown as a function of time. These data are replotted as ratios relative to PC (red line, set to 2 to reflect the bilayer composition ratio) in **D**.

### Interactions in atomistic simulations correlate with experimental observations

To explore the interactions predicted by CG simulation in more detail alongside possible conformational changes on binding to the membrane, we performed atomistic (AT) simulations starting from instances of α-synuclein binding to the membrane taken from the CG/POPG simulations. Thus, we selected 3 representative structures of the bound monomer from the CG simulations and converted them to AT resolution (44). Each of these 3 systems formed the starting point for 10 replicate AT-MD simulation each of duration 0.1 to 0.25 μs, yielding an aggregated simulation time of 5.5 μs (see Fig. 3 and Table 1).

**Figure 3:**
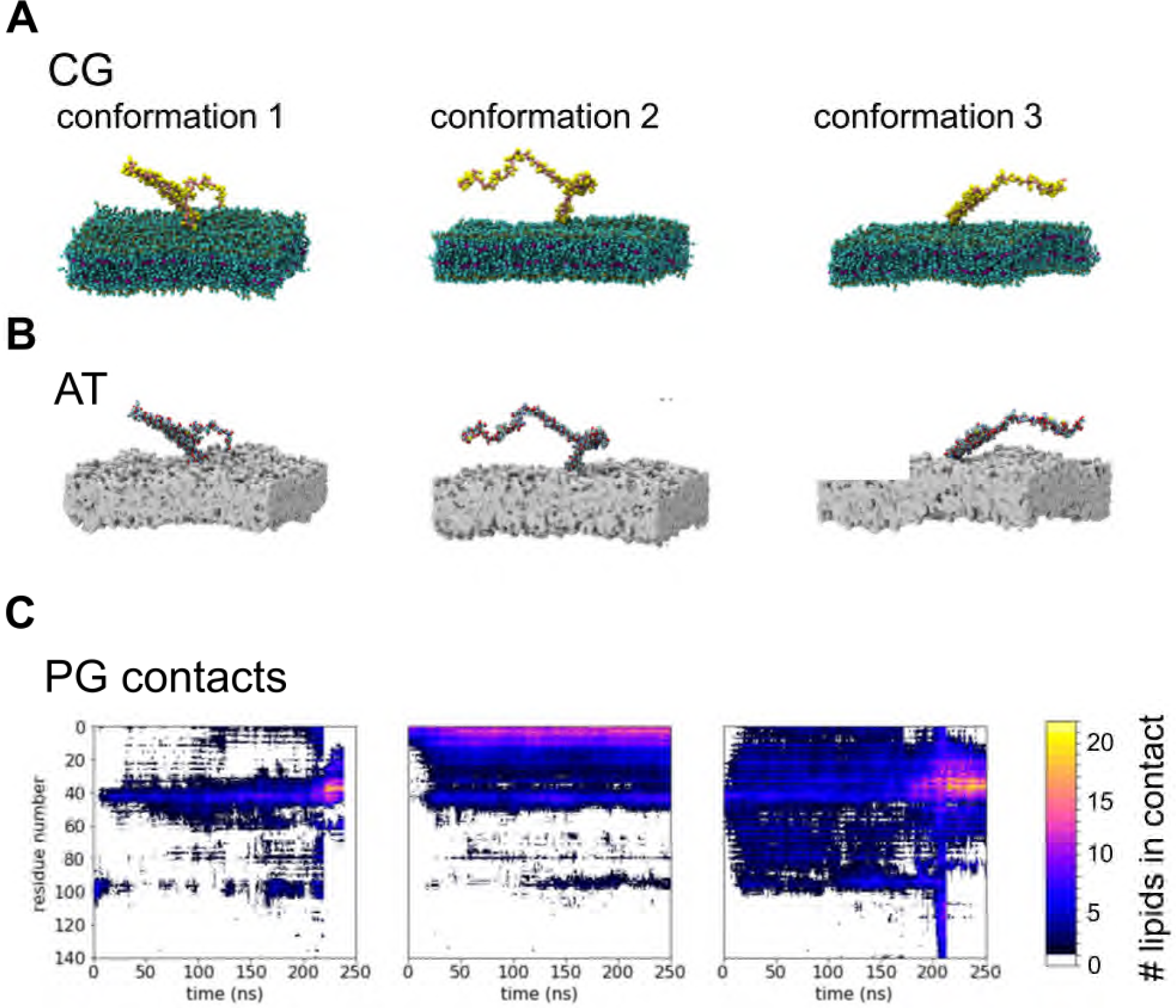
Atomistic simulations of the interaction of α-synuclein with a POPG membrane. **A** Three binding poses from the CG simulations are shown, they were chosen to capture initial interactions at the C-terminus (1 - left), at the N-terminus (2-centre) and at the inter-helical region (3-right). These were converted to the corresponding atomistic models shown in **B** to initiate atomistic simulations (of duration 100-250 ns with 10 replicates), resulting in an aggregated simulation time of 5.5 *μ*s. **C** The number of contacts to lipids within a 1 nm cutoff of the Cα atom of each residue is shown as a function of time and residue number for each starting model, averaged across replicates.

Analysis of the lipid contacts as a function of time (Fig. 3C) shows the interhelical region generally binds first, with the disordered C-terminus making transient contacts with the membrane (Fig. 3C). Where the C-terminus is initially in contact, this is followed by reorientation of the protein to bring the interhelical region closer to the membrane. Whilst both the contact profile and the bilayer penetration are variable across the ensemble of simulations, overall it appears that the α-synuclein monomer prefers to bind initially through the interhelical region, but that there is some flexibility in the binding pose.

During the course of the AT simulations we frequently observed a break in the middle of Helix 2. A representative trajectory (Fig. 4A) shows Helix 2 (residues 46-95) breaking between residues 65-70 from about 40 ns onwards. Averaging over all 10 replicate simulations (Fig. 4B) shows that residues 60-70 have a substantially reduced probability of adopting an α-helical conformation. This profile is in excellent agreement with the equivalent profile derived from NMR chemical shift data for a-synuclein bound to bicelles (DHPC:DMPG:PIP_2_ ∼3:5:1) (Fig. 4C; also see SI Fig. S3), which shows reduced α-helicity in the same region, with values nearly as low as in the inter-helical region. The regions of high probability of forming α-helical structure observed in computational studies matches well with the regions of higher α-helical propensity as observed in NMR-measurements. The small differences in the profiles may reflect the different lipids employed (POPG vs. DHPC:DMPG:PIP_2_), although in both the simulations and the experiments the membrane surface was predominantly anionic.

**Figure 4:**
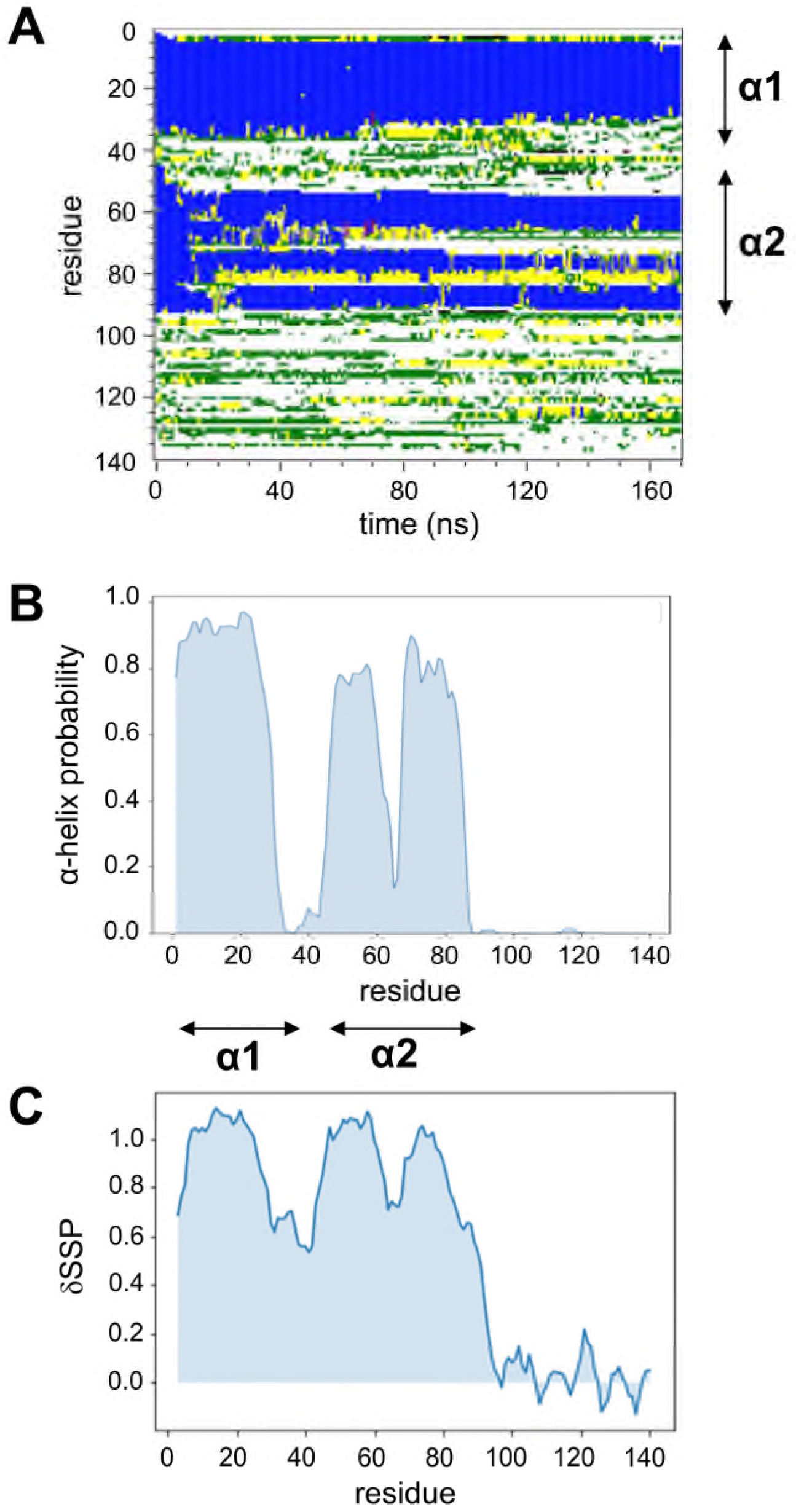
Secondary structure of α-synuclein, comparing atomistic simulations when bound to a POPG bilayer with NMR data. **A** Secondary structure (as defined by DSSP) as a function of residue number and time for a representative trajectory corresponding to conformation 1 (see Fig. 3 above). Loss of helical structure in the middle of helix 2 (in the region of residue 70) is observed. The locations of helices 1 and 2 in the NMR structure (PDB id 1XQ8) are indicated to the right of the diagram. **B** Frequency of α-helical secondary structure as a function of residue averaged across all simulations of conformation 1. The fraction of α-helix is reduced in favour of random coil conformations between residues 60-70. (Similar profiles are seen for simulations of conformations 2 and 3). **C** NMR Chemical shift data showing secondary structure propensities. Chemical shift indexing shows field-shifted atoms in the region between residues 60-75, indicating a reduction in propensity of α-helical structure. The values obtained are the average of the shift observed versus random coil expected shifts, weighted by their sensitivity to α-helical or extended conformations. Data are shown for α-synuclein bound to bicelles composed of DHPC, DMPG and PIP_2_ (see Methods for details). The extent of the two helices in the SDS-bound structure (PDB id 1XQ8) are again indicated by arrows.

In a number of the AT simulation trajectories β-sheet structure is observed in some residues, mostly in the middle of Helix 2 and around the interhelical region (SI Fig. S4) Taken with the CG results, which show Helix 2 to preferentially sit on top of membrane bound Helix 1, this might suggest that extended β-structures could be seeded in the inter-helical region or NAC region, thereby initiating hydrophobic aggregation of α-synuclein molecules.

We have also compared our simulations of α-synuclein at the bilayer surface with experimental data on the larger scale structure of the protein, obtained via cross-linked mass spectrometry (XLMS) studies (Fig. 5). The large number of different crosslinks observed (see SI Fig. S5) is as expected for the highly dynamic conformational ensemble of the IDP α-synuclein, which retains a large degree of flexibility even in its membrane-bound state (45). The XLMS-data shown presents the sum of all peptide spectrum matches (PSMs) found between two positions of the protein as 2D “contact map”. As X-linking positions are restricted to the N-terminus, Lys, Asp and Glu residues and the amount of PSMs found vary strongly, we applied a density estimation to the PSM pattern found for easier visual comparison of XLMS and simulation data. (The underlying contact map is shown in SI Fig. S5A) Comparison of the XLMS data and an intra-monomer contact matrix derived from a simulation trajectory (Fig. 5) shows good agreement between experimental measurements and the simulation. This is especially true regarding the off diagonal (OD) contacts observed between regions 50-70 and 10-30. The full diagonal visible in the simulation data (Fig 5B) is missing from the XLMS picture due to chemical crosslinker used, the restrictions in places on the amino acids to be linked and the requirement for the PSMs to stem from two separate peptides. Interestingly the N-terminal region that shows many PSMs close to the diagonal is dominated by PSMs with an amino acid spacing of 3, 4 or 6 residues fitting with the assumption that they link side-chains within an α-helix. The observed strong diagonal (SD) caused by a higher amount of PSMs at the N-terminus fit with NMR and simulation observations of a stable Helix 1 at the N-terminus when the protein is membrane bound (45, 46).

**Figure 5:**
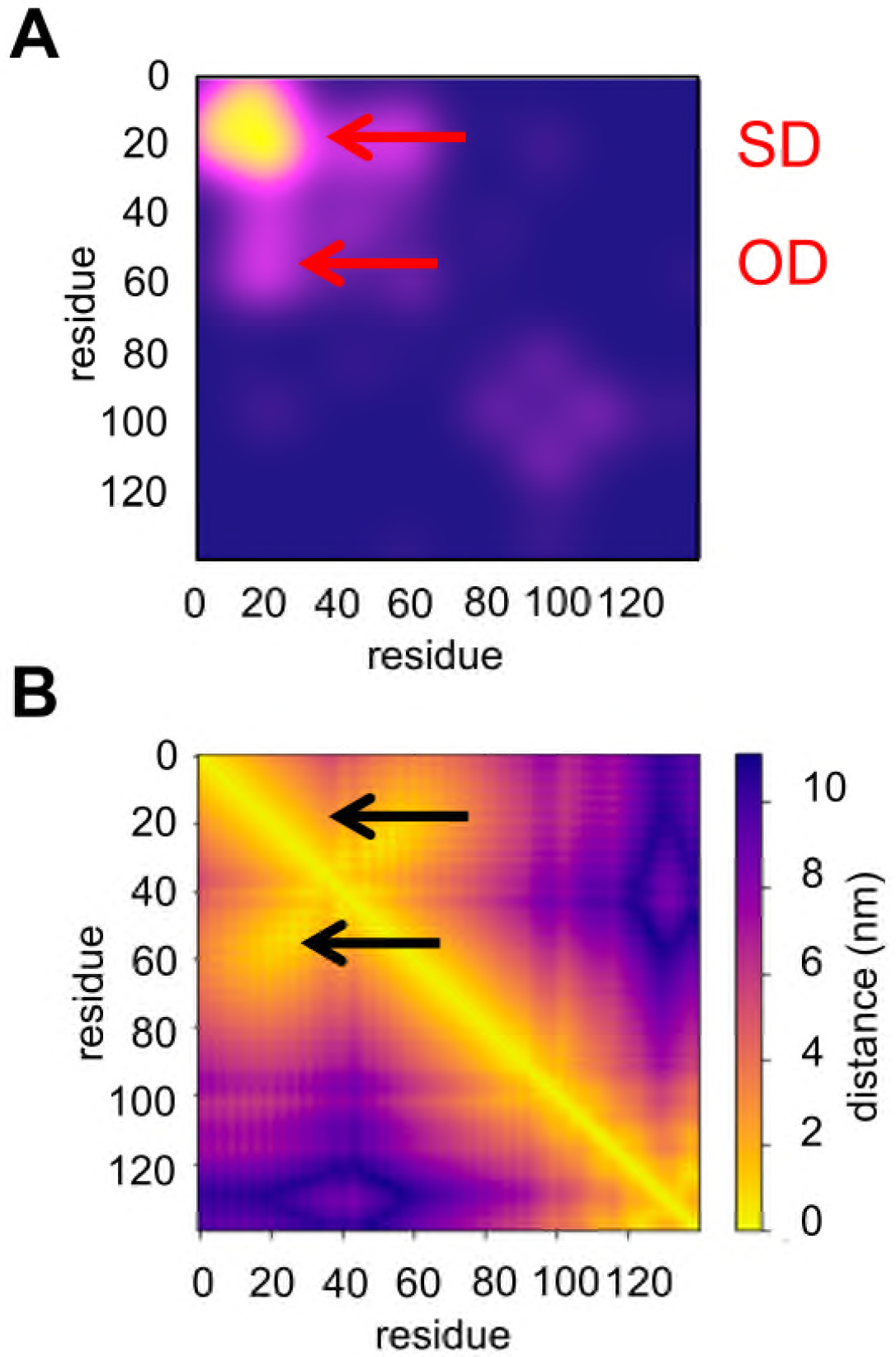
Residue-residue distances of the α-synuclein monomer binding to a POPG bilayer. **A** Density estimation of the XLMS Crosslink-Pattern observed in α-synuclein when bound to POPG liposomes (based on PSM counts for each residue pair). **B** Residue-residue distances generated from the simulation ensemble from conformation 2. (Similar profiles are seen for simulations of conformations 1 and 3). Arrows indicate a strong peak close to the diagonal (SD) corresponding to links driven likely by a helical conformation and an off-diagonal peak (OD) which is seen clearly in both **A** and **B** and which corresponds to contacts between helix 1 and the N-terminal segment of helix 2.

Although it is not as disordered as in solution, α-synuclein retains a high degree of flexibility in its membrane bound state (45). As noted above, this leads to a large number of detected crosslinks and their spatial distribution precludes us from evaluating the crosslinks found on the basis of a single structure. Recently studies on the solution state of α-synuclein have used a large data set generated with different cross-linkers as the basis for constraint-guided discrete MD in order to calculate an ensemble of structures fulfilling XLMS derived distance restraints (12). Although the membrane bound state of α-synuclein is less flexible, both the lower amount of distance information available as well as the poor matching of NMR and SAXS derived ensembles with those generated using XLMS data led us to use a different approach. Here we show the similarity in interacting regions as derived from experimental and computational approaches. Both of these capture a large ensemble of structures and without explicit calculation of ensembles for the experimental data we can demonstrate similar behaviour as observed in simulations. The SI Fig. S5C demonstrates the intramolecular contacts within the bound α-synuclein as analysed by AT simulations, which are represented as heat maps. In all simulations, the off-diagonal shows that Helix 1 and Helix 2 lie adjacent to each other. As the XLMS data stems from the bound form of the protein, where long contact times lead to strong binding of the N-terminus of the protein (45) this observation matches with the observed best agreement between XLMS data and the profile of conformation 2.

### Markov state models of the conformational changes of membrane-bound α-synuclein

To further investigate the conformational changes of the α-synuclein monomer when bound to a POPG bilayer, we constructed Markov state models using MSMBuilder (47). In these we focussed on the conformational dynamics of residues 60-70 of the bound α-synuclein monomer as our previous analysis (above; Fig. 4) had indicated that this is where the break in Helix 2 occurs. We therefore used the pooled data from the AT-MD simulations to construct an MSM featurised on the Cα contacts of these. We plotted the implied timescales to check for convergence and chose an MSM lag time of 2 ns for model construction (Fig. 6A). Fig.6B shows the clustered states coloured by the first eigenvector, which represents the slowest transition from red states to blue. Finally we clustered these microstates into six macrostates. An appropriate number of macrostates is one more than the number of timescales above the major gap (48). Thus, for our system the timescales would indicate choosing either two or three. However, the energy landscape indicates there may be more than three (in particular the positioning of microstates in Fig.6B) and it has been recommended to err on the side of having too many microstates rather than too few. We generated representative coordinates from the two lowest energy states (see Fig. 6C) and visualised them (Fig. 6DE). It can be seen that the MSM captures a conformational transition from an α-helical structure for residues 60-70 to a conformation in which there is local loss of helicity with the peptide chain folded back on itself. A conformational change in this region is demonstrated in line with the NMR secondary shift propensity data.

**Figure 6:**
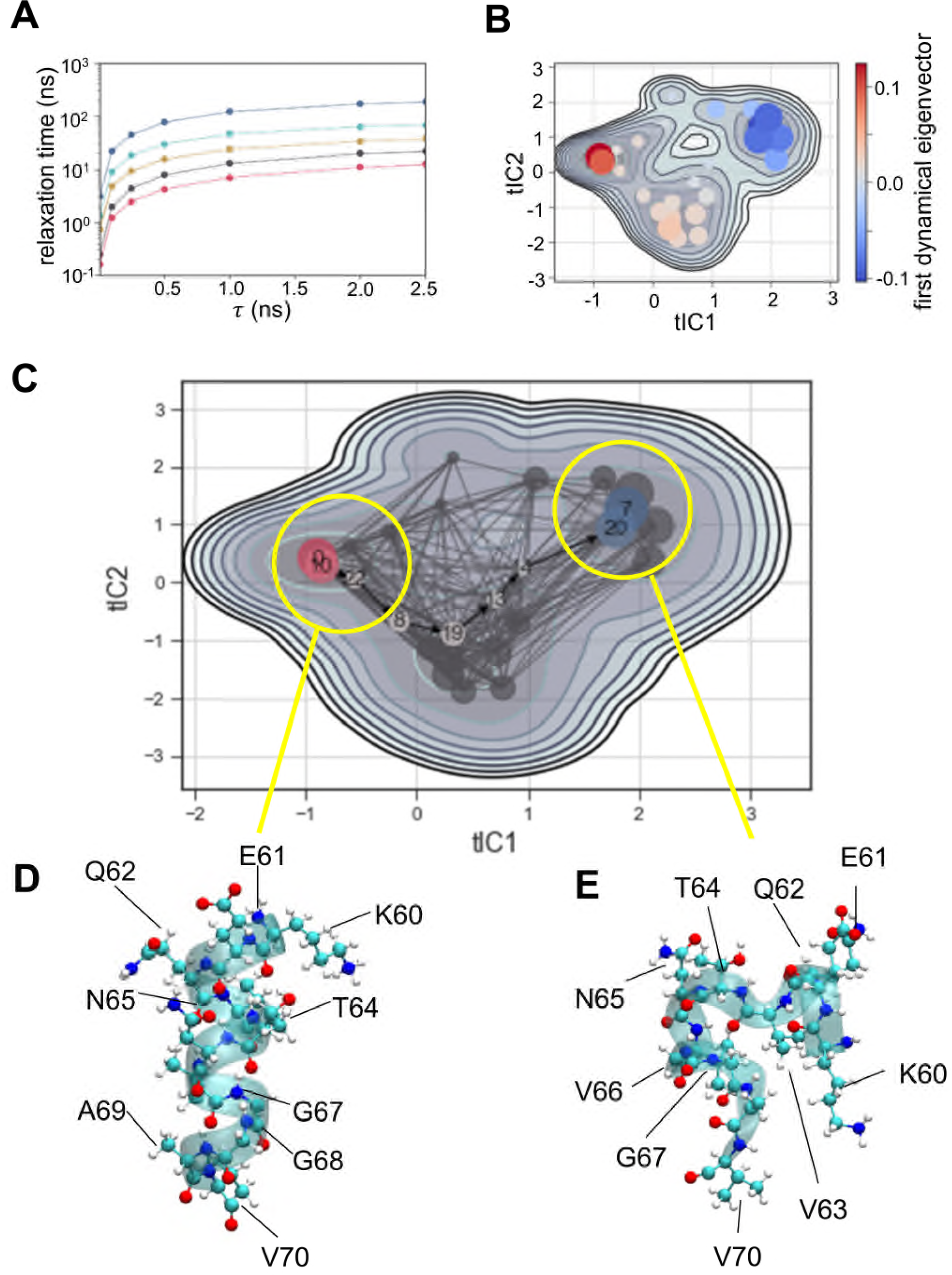
Markov state model of residues 60-70. **A** Implied timescales are plotted to choose the lag time for the MSM. Implied timescales show processes converge at a lag time of 200 frames (2ns). **B** Slowest motion as represented by the 1st dynamical eigenvector from red states to blue. This corresponds to the slowest structural motion observed across the ensemble and can be interpreted as the major structural change. **C** MSM microstate transitions and representative structures. Yellow circles indicate the two lowest energy macrostate centres from which representative structures (**D, E**) were extracted. Thus **D** shows the α-helical structure of residues 60-70. This is generated from a region that is structurally close to the conformation at the start of the trajectory. **E** shows a break in the helix, such that the structure around residues 60-70 turns back on itself.

This local structure with a bend around residue V66 (Fig. 6E) is of interest because amyloid structures for α-synuclein determined by cryo-EM (49, 50) show a kernel structure near the dimer interface which also has a bend around V66. Our simulations suggest an attractive hypothesis that the initial bent conformation formed on the bilayer surface may be a potential seed for subsequent amyloid formation. A small molecule thought to interact with residues around 53-73, over the region to which we have built the model, inhibits in vitro aggregation of α-synuclein (51).

## Conclusions

We have performed a multiscale MD simulation study to investigate the membrane-binding mechanism of α-synuclein. The initial CG simulations allow us to propose the major contacts between the protein and an anionic lipid bilayer emphasizing the role of the interhelical loop region in initiating membrane binding. Subsequent atomistic simulations suggest the formation of a non-helical region in the centre of Helix 2 following membrane binding. This model is in agreement with NMR chemical shift data for α-synuclein bound to anionic lipid bicelles. Additionally, convincing agreement was also found for Helix 1-Helix 2 contacts as identified by MD simulations as well as chemical cross-linking mass spectrometry. Markov state models suggest that the non-helical region enables a conformational transition which forms a bend in the centre of the Helix 2 region, which in turn may correlate with a key bend in the proposed structure of the amyloid fibril formed by α-synuclein.

## Methods

### Computational

The NMR structure of micelle-bound α-synuclein (17) (PDB id 1XQ8) was used and converted to a CG representation. CG simulations of α-synuclein were performed using a 10 × 10 nm^2^ area bilayer of POPG or a lipid mixture of DOPC/DOPE/DOPS in a 2/5/3 ratio (see Table 1). Bilayers were built using INSANE (52). The protein molecule was positioned 4 nm away from the bilayer surface. The box was solvated and sodium and chloride ions added to a concentration of ∼0.15 M. CG simulations were performed using GROMACS 5.1 (53, 54). Energy minimisation was carried out via steepest descent and the system equilibrated for 5 ns with protein backbone particles restrained. Productions simulations were run without restraints for 1 or 2 μs with 10 replicates with different initial velocities (Table 1).

CG simulations were performed using the Martini 2.1 force field (55)with a 20 fs time step. Particle coordinates were written out every 0.5 ns. Coulombic interactions were shifted to zero between 0 and 1.2 nm. Lennard-Jones interactions were shifted to zero between 0.9 and 1.2 nm. The nearest neighbour list was updated every 10 steps. The Berendsen thermostat (56) (coupling constant 1 ps) and barostat (coupling constant 1 ps, compressibility 5×10^−6^ bar^-1^) were used to maintain temperature at 323 K and pressure at 1bar. The LINCS algorithm (57) was used to constrain bond lengths.

AT simulations were performed using the Charmm36 force field (58) with a 2 fs time step. Atomic coordinates were written out every 20 ps. Lennard-Jones interactions were shifted to zero between 0.9 and 1.2 nm. Long-range electrostatic interactions were treated using the particle-mesh Ewald method (PME) (59) using default parameters pme-order = 4 and ewald-rtol = 10^−5^, fourierspacing = 0.12. PME was shifted from 0 to 1 nm (40). The nearest neighbour list was updated every 10 steps. A velocity-rescale thermostat (60) (coupling constant 1 ps) and Parrinello-Rahma (61) barostat (coupling constant 1 ps, compressibility 5×10^−6^ bar^-1^) were used to maintain the temperature and pressure. The LINCS algorithm was used to constrain bond lengths.

AT simulations were started from snapshot structures taken from CG simulation and converted to atomistic representations (62). AT-MD simulations were performed using GROMACS 5.1/2018. The system was equilibrated for 1.5 ns with the backbone atoms of the protein restrained, then a production run was performed (Table 1).

VMD was used for simulation visualisation (63). Graphs were generated in matplotlib (64) and seaborn. Contact analysis was carried out with in house python scripts using a cutoff of 0.7 nm for CG and 1.0 nm for atomistic from the Cα carbon atom. PCA and correlation analysis were carried out using standard libraries in matplotlib/seaborn and scikit-learn (65). Markov state modelling was carried out in MSMBuilder (47). The system was featurised on backbone contacts and scaled using StandardScaler. The number of microstates was set at 50 for calculation and implied timescales plotted to assess convergence. The system was clustered into macrostates (this is further explained in the context of the results) and the two lowest energy states were used to extract representative structures which were visualised with VMD.

### Experimental

#### NMR

α-Synuclein was expressed and purified as described previously (66). Secondary shift propensity scores (SSPs) were calculated from H, HN, CO, Cα and Cβ shift values obtained in phosphate buffer pH = 5.5, 323K. They were calculated using the SSP-script (67) using Cα and Cβ to apply internal referencing. Assignments of α-synuclein in these conditions were obtained using TROSY (68) versions of 3D backbone assignment pulse sequences for the bicelle bound form of α-synuclein. In order to obtain sufficient signal intensity the protein was partially deuterated by expression in D_2_O based M9 media. Bicelles were composed of DHPC, DMPG and PiP_2_ with final concentrations of 100µM α-synuclein, 10.6mM DHPC, 5.1mM DMPG and 1.1mM PIP_2_. As a significant fraction of DHPC is expected to from Micelles, the Bicelles are expected contain about 3.6mM DHPC.

#### XLMS

Crosslinking of α-synuclein was carried out with the zero-length 1-ethyl-3-(3-dimethylaminopropyl)carbodiimide (EDC) in the presence of liposomes composed of POPG in phosphate buffer pH=6.5. 50µM of α-synuclein was incubated for 60 min at room temperature in the dark with 2mM EDC and 5mM Hydroxy-2,5-dioxopyrrolidine-3-sulfonicacid (Sulfo-NHS) with 1mg/ml of POPG-based liposomes present during the reaction. Under these conditions the majority of α-synuclein is bound to the liposomal surface (66). The reaction was stopped with Tris (50mM) and β-mercaptoethanol (20mM). The resulting crosslinked protein was then subjected to SDS-PAGE, which showed monomer, dimer and multimer bands of α-synuclein. The band corresponding to the monomeric weight of α-synuclein was selected for further analysis in order to avoid interference by crosslinks stemming from intermolecular interactions. Liposomes used in this reaction were produced as described previously (66).

The monomer band was excised from the gel and destained with a mixture of acetonitrile (Chromasolv®, Sigma-Aldrich) and 50 mM ammonium bicarbonate (Sigma-Aldrich). The proteins were reduced using 10 mM dithiothreitol (Roche) and alkylated with 50 mM iodoacetamide. Trypsin (Promega; Trypsin Gold, Mass Spectrometry Grade) and chymotrypsin (Promega sequencing grade) were used for proteolytic cleavage. Digestion was carried out with trypsin at 37°C overnight and subsequently with chymotrypsin at 25°C for 5h. Formic acid was used to stop the digestion and extracted peptides were desalted using C18 Stagetips (69).

Peptides were analysed on an UltiMate 3000 HPLC RSLC nano system (Thermo Fisher Scientific) coupled to a Q Exactive HF mass spectrometer (Thermo Fisher Scientific), equipped with a Nanospray Flex ion source (Thermo Fisher Scientific). The samples were loaded on a trap column (Thermo Fisher Scientific, PepMap C18, 5 mm × 300 μm ID, 5 μm particles, 100 Å pore size) at a flow rate of 25 μL min^-1^ using 0.1% TFA as mobile phase. After 10 min, the trap column was switched in-line with the analytical C18 column (Thermo Fisher Scientific, PepMap C18, 500 mm × 75 μm ID, 2 μm, 100 Å) and peptides were eluted applying a segmented linear gradient from 2% to 80% solvent B (80% acetonitrile, 0.1% formic acid; solvent A 0.1% formic acid) at a flow rate of 230 nL/min over 120 min. The mass spectrometer was operated in data-dependent mode, survey scans were obtained in a mass range of 350-1650 m/z with lock mass activated, at a resolution of 120,000 at 200 m/z and an AGC target value of 3E6. The 10 most intense ions were selected with an isolation width of 1.6 Thomson for a max. of 250 ms, fragmented in the HCD cell at 28% collision energy and the spectra recorded at a target value of 1E4 and a resolution of 60,000. Peptides with a charge of +1,+2 or >+7 were excluded from fragmentation, the peptide match feature was set to preferred, the exclude isotope feature was enabled, and selected precursors were dynamically excluded from repeated sampling for 20 seconds within a mass tolerance of 8 ppm.

For peptide and protein identification raw data were processed using the MaxQuant software package (70) (version 1.5.5.1) and spectra searched against a combined database of the alpha-synuclein construct sequence, the E.coli K12 reference proteome (Uniprot) and a database containing common contaminants. The search was performed with full trypsin and chymotrypsin specificity and a maximum of three missed cleavages at a protein and peptide spectrum match false discovery rate of 1%. Carbamidomethylation of cysteine residues were set as fixed, oxidation of methionine and N-terminal acetylation as variable modifications. All other parameters were left at default. To identify cross-linked peptides the spectra were searched using pLink (71) (version 1.23). Q Exactive HF raw-files were pre-processed and converted to mgf-files using pParse (72). The spectra were searched against a database containing the 8 most abundant protein hits (sorted by MS/MS counts) identified in the MaxQuant search. Carbamidomethylation of cysteine and oxidation of methionine residues were set as variable modifications. Trypsin/Chymotrypsin was set as enzyme specificity, EDC was set as cross-linking chemistry allowing Asp and Glu residues being linked to Lys residues. Search results were filtered for 1% FDR at the PSM level and a maximum allowed precursor mass deviation of 5 ppm. To remove low quality peptide spectrum matches an additional e-Value cutoff of < 0.001 was applied. Proteomics analyses were performed by the Max Perutz Laboratories Mass Spectrometry Facility using the VBCF instrument pool.

The resulting pattern of crosslinks is very varied due to the high flexibility of bound α-Synuclein. Looplinks were not included in this analysis as these would show short range contacts only. We show the resulting crosslink-pattern as a density plot generated with the 2D kernel density estimation using stat_density2d in R with parameter geom="tile” generated with the R-package ggplot2 (73) (Figure 5A).

## Acknowledgements

The authors would like to thank A.L. Biere and the UCB α-synuclein teams for strategic discussions. Research in M.S.P.S.’s group is funded by BBSRC, EPSRC (ARCHER), EU (PRACE), and the Wellcome Trust. We thank UCB for additional support.

## Supporting Information

**SI Figure S1:**
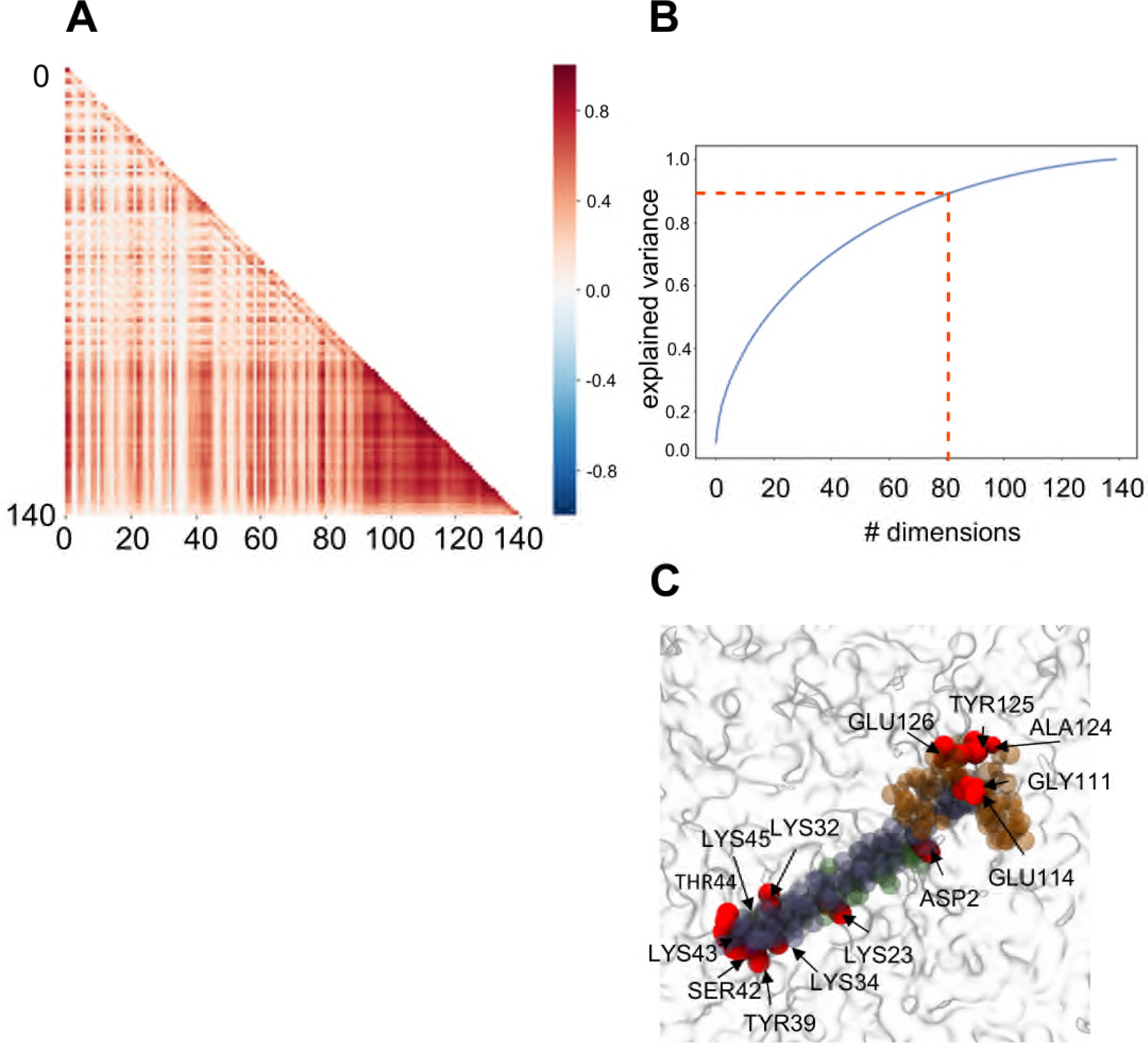
Analysis of coarse grained simulations of the interaction of α-synuclein with a PG bilayer. **A** Correlation analysis of protein residue-PG contacts, showing positive correlation in contacts between the disordered tail and the interhelical region and residues 79-80. **B** Principal components analysis showing that 80 residues explain 95% of the variance in binding. **C** View from above of a representative PG-bound structure of α-synuclein with the top 14 residues making lipid contacts highlighted in red (see also Fig. 1E). PG headgroups are shown in grey.

**SI Figure S2:**
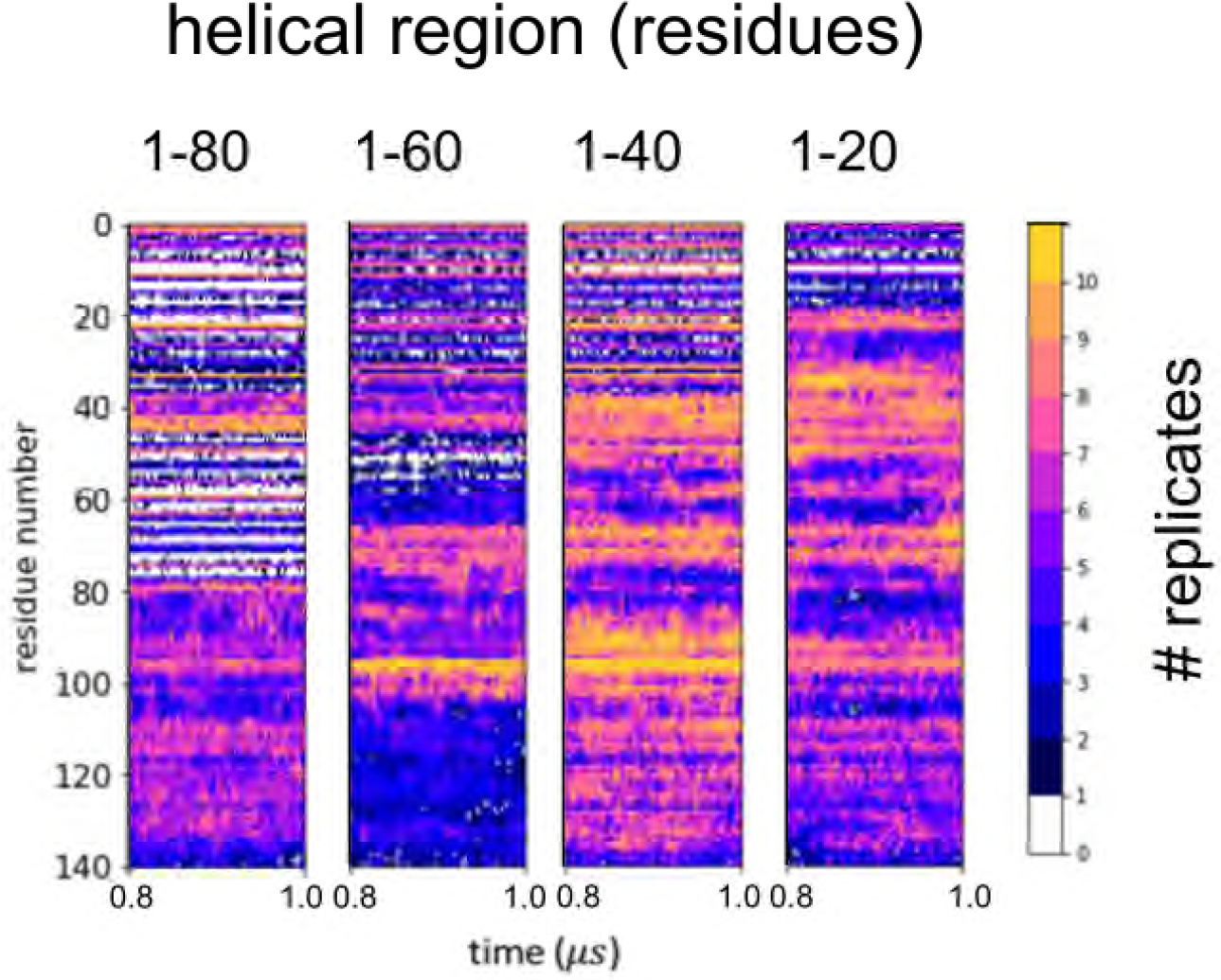
Coarse grained simulations of the interactions with a PG membrane of α-synuclein models with varying degrees of helicity. Each conformation was simulated for 10 replicates of 1 μs. The colours indicate the number of replicates across the ensemble making contacts at that time point. Only the last 0.2 μs of each set of simulations are shown. The contact profiles show that the contribution from the inter-helical region remains a common feature of binding, but abrogation of helicity results in a greater contribution from residues 45-100.

**SI Figure S3:**
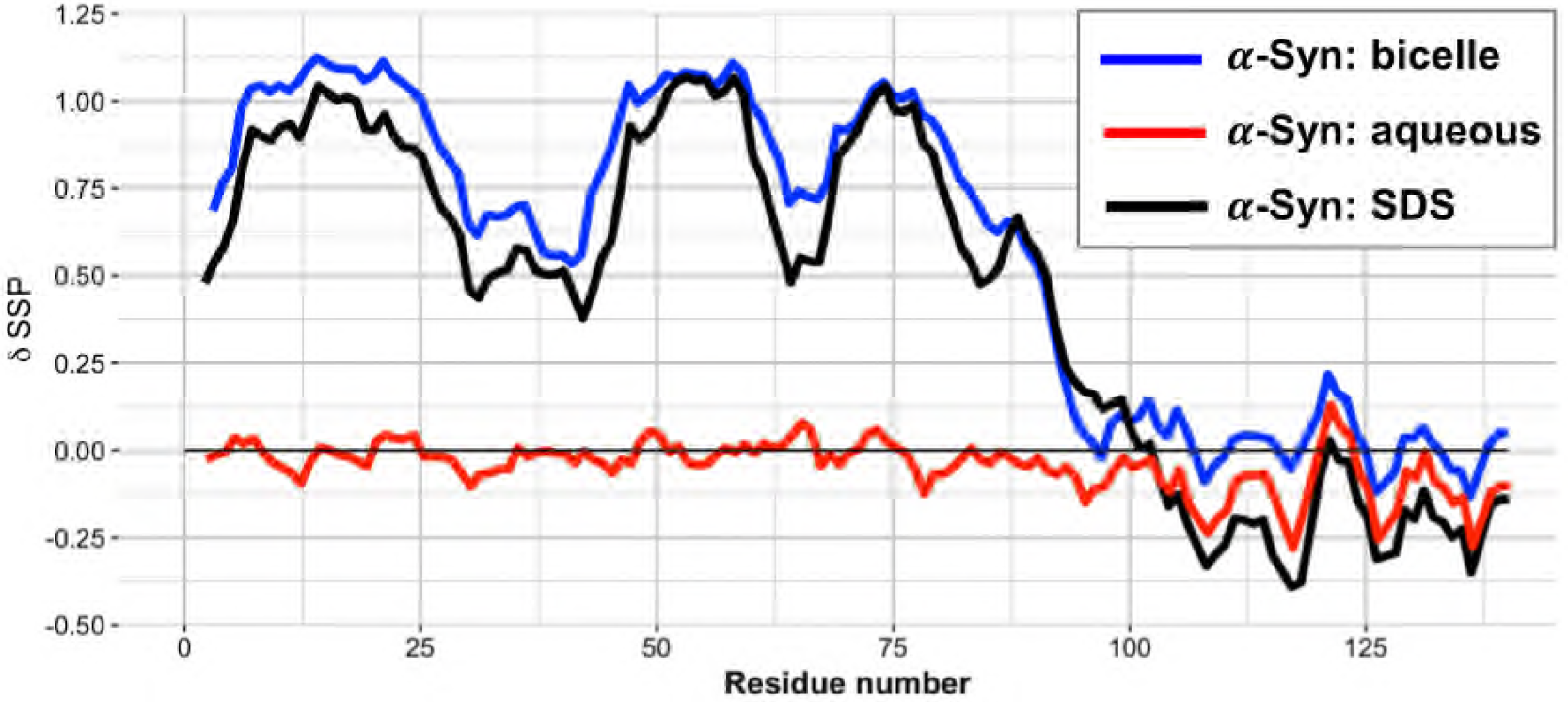
Comparison of secondary structure propensity (SSP) scores calculated from NMR measurements from samples α-synuclein bound to lipid bicelles (blue; see main text, Figure 4C), to SDS micelles (black) or in aqueous solution (red). All were assigned at pH=5.5 and measured at 323 K. This shows that changes with SDS and Bicelles are similar and that the free ensemble has no clearly distinguishable secondary structure. As described in the main text, secondary structure propensity values were calculated from Cα, Cβ, NH, N and CO values using the program SSP. The values obtained are the average of the shift observed versus random coil expected shifts, weighted by their sensitivity to alpha-helical or extended conformations. The observed Cα and Cβ shifts were also used for shift referencing, which removes the pH-dependent effects almost entirely. All three SSP score series were determined from α-synuclein samples recorded at the same buffer and temperature conditions. Assignments of the above conditions were carried out using standard-triple resonance experiments and include the used side-chain resonances, but do not include side-chain assignments.

**SI Figure S4:**
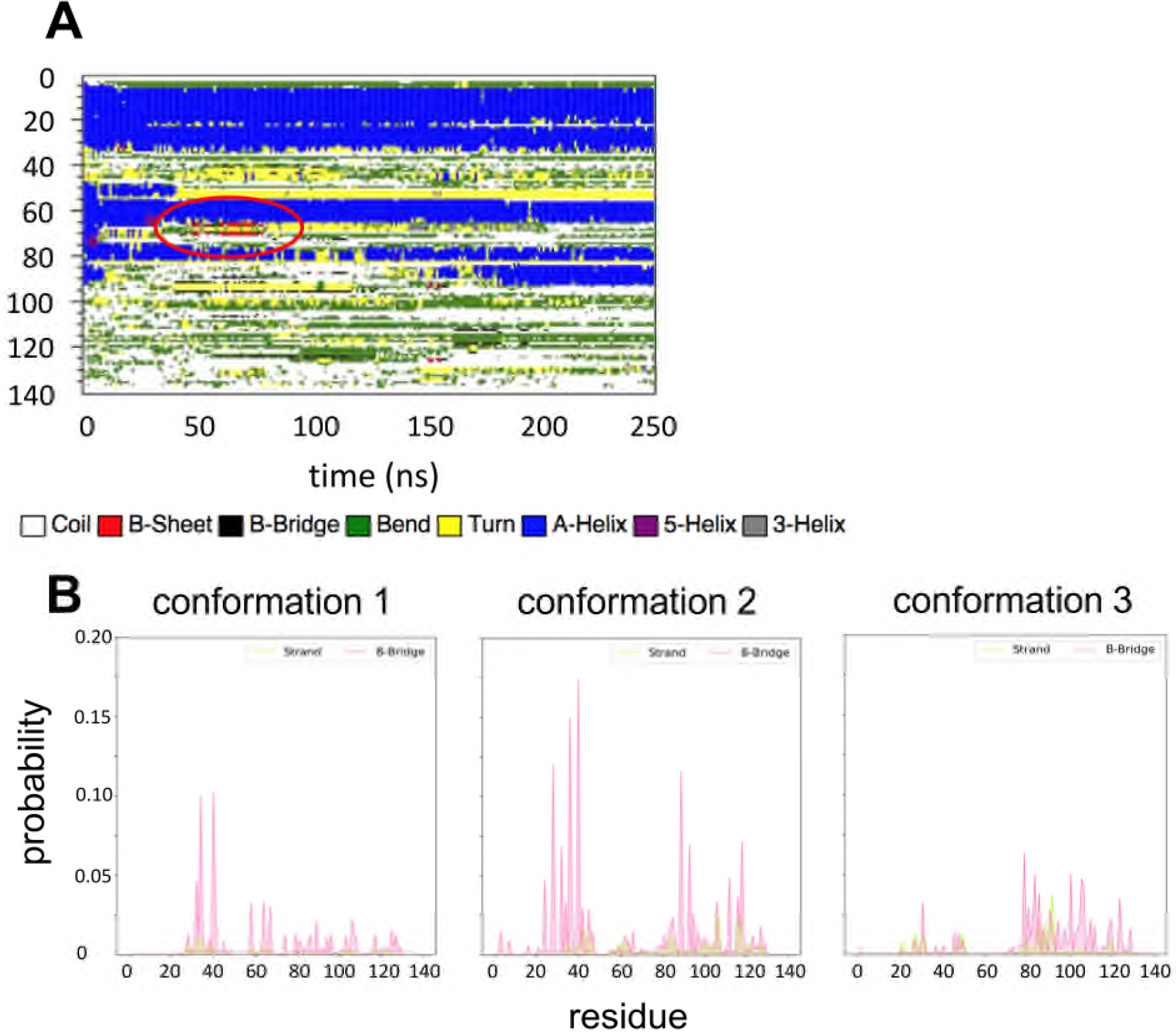
Secondary structure (DSSP) analysis of atomistic simulations of α-synuclein binding to a POPG membrane. **A** DSSP analysis for one trajectory from starting conformation 2 (see main text, Fig. 3 for details). The trajectories show β-sheet structure emerging (highlighted by a red ellipse) around the interhelical region and in the NAC region. **B** Probability of β-strand and β-bridge assignment for each of three atomistic simulations starting from different conformations (see main text, Fig. 3 for details).

**SI Figure S5:**
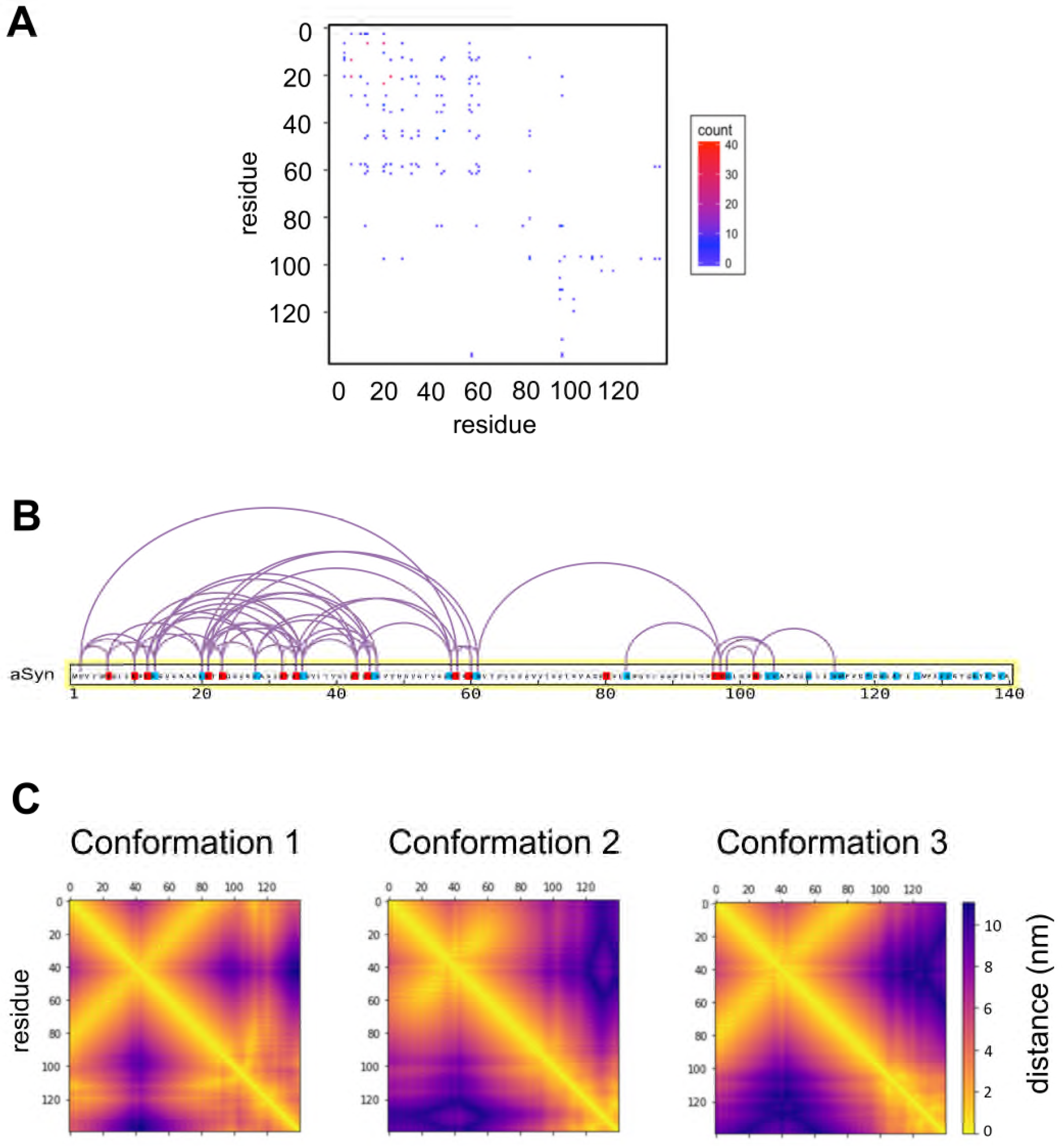
Results of cross-linked mass spectrometry (XLMS) studies and comparison with contacts in simulations (see main text and Fig. 5 for details). The XLMS-data are used to determine a sum of all peptide spectrum matches (PSMs) found between two positions of the protein. This is shown in **A** as a pattern of PSMs determined with *binwidth* = 1, indicating the number of PSMs for each individual crosslink pair. This corresponds to the crosslinks and provides the data used in all subsequent plots (see Fig. 5A). **B** shows the XLMS-pattern mapped onto the sequence of α-synuclein with K (blue), E and D (red) residues indicated. **C** Residue-residue distances generated from the atomistic simulation ensembles from conformation 1, 2 and 3 (see also main text Fig. 5B).

## References

1. Goedert M, Jakes R, Spillantini MG (2017) The synucleinopathies: Twenty years on J Parkinsons Dis 7:S53–S71.

2. Polymeropoulos MH, et al. (1997) Mutation in the alpha-synuclein gene identified in families with Parkinson’s disease Science 276:2045–2047.

3. Burre J, et al. (2010) α-Synuclein promotes SNARE-complex assembly in vivo and in vitro Science 329:1663–1667.

4. Oliveira LMA, et al. (2015) Elevated α-synuclein caused by SNCA gene triplication impairs neuronal differentiation and maturation in Parkinson’s patient-derived induced pluripotent stem cells Cell Death & Dis 6: e1994.

5. Jin HJ, et al. (2011) alpha-Synuclein negatively regulates protein kinase Cd expression to suppress apoptosis in dopaminergic neurons by reducing p300 histone acetyltransferase activity J Neurosci 31:2035–2051.

6. Ugalde CL, Finkelstein DI, Lawson VA, Hill AF (2016) Pathogenic mechanisms of prion protein, amyloid-β and α-synuclein misfolding: the prion concept and neurotoxicity of protein oligomers J Neurochem 139:162–180.

7. Weinreb PH, et al. (1996) NACP, a protein implicated in Alzheimer’s disease and learning, is natively unfolded Biochemistry 35:13709–13715.

8. Fauvet B, et al. (2012) α-Synuclein in central nervous system and from erythrocytes, mammalian cells, and escherichia coli exists predominantly as disordered monomer J Biol Chem 287:15345–15364.

9. Theillet FX, et al. (2016) Structural disorder of monomeric α-synuclein persists in mammalian cells Nature 530:45–50.

10. Schwalbe M, et al. (2014) Predictive atomic resolution descriptions of intrinsically disordered hTau40 and α-Synuclein in solution from NMR and small angle scattering Structure 22:238–249.

11. Allison JR, Varnai P, Dobson CM, Vendruscolo M (2009) Determination of the free energy landscape of α-synuclein using spin label nuclear magnetic resonance measurements J Amer Chem Soc 131:18314–18326.

12. Brodie NI, et al. (2019) Conformational ensemble of native α-synuclein in solution as determined by short-distance crosslinking constraint-guided discrete molecular dynamics simulations PLoS Comput Biol 15: e1006859.

13. Brodie NI, Petrotchenko EV, Borchers CH (2016) The novel isotopically coded short-range photo-reactive crosslinker 2,4,6-triazido-1,3,5-triazine (TATA) for studying protein structures J Proteomics 149:69–76.

14. Uluca B, et al. (2018) DNP-Enhanced MAS NMR: A tool to snapshot conformational ensembles of α-synuclein in different states Biophys J 114:1614–1623.

15. van Maarschalkerweerd A, et al. (2014) Protein/lipid coaggregates are formed during α-synuclein-induced disruption of lipid bilayers Biomacromolecules 15:3643–3654.

16. Drescher M, et al. (2008) Antiparallel arrangement of the helices of vesicle-bound α-synuclein J Amer Chem Soc 130:7796–7797.

17. Ulmer TS, Bax A, Cole NB, Nussbaum RL (2005) Structure and dynamics of micelle-bound human alpha-synuclein J Biol Chem 280:9595–9603.

18. Trexler AJ, Rhoades E (2009) α-synuclein binds large unilamellar vesicles as an extended helix Biochemistry 48:2304–2306.

19. Bartels T, Choi JG, Selkoe DJ (2011) α-synuclein occurs physiologically as a helically folded tetramer that resists aggregation Nature 477:107–U123.

20. Ulmer TS, Bax A (2005) Comparison of structure and dynamics of micelle-bound human alpha-synuclein and Parkinson disease variants J Biol Chem 280:43179–43187.

21. Sode K, Ochiai S, Kobayashi N, Usuzaka E (2007) Effect of reparation of repeat sequences in the human α-synuclein on fibrillation ability Int J Biol Sci 3:1–7.

22. Flagmeier P, et al. (2016) Mutations associated with familial Parkinson’s disease alter the initiation and amplification steps of α-synuclein aggregation Proc Natl Acad Sci USA 113:10328–10333.

23. Bengoa-Vergniory N, Roberts RF, Wade-Martins R, Alegre-Abarrategui J (2017) α-synuclein oligomers: a new hope Acta Neuropathol 134:819–838.

24. Winner B, et al. (2011) In vivo demonstration that α-synuclein oligomers are toxic Proc Natl Acad Sci USA 108:4194–4199.

25. Chen SW, et al. (2015) Structural characterization of toxic oligomers that are kinetically trapped during α-synuclein fibril formation Proc Natl Acad Sci USA 112:E1994–E2003.

26. Outeiro TF, et al. (2008) Formation of toxic oligomeric α-synuclein species in living cells PLoS One 3:e1867

27. Pineda A, Burre J (2017) Modulating membrane binding of α-synuclein as a therapeutic strategy Proc Natl Acad Sci USA 114:1223–1225.

28. Graen T, et al. (2018) Transient secondary and tertiary structure formation kinetics in the intrinsically disordered state of α-synuclein from atomistic simulations ChemPhysChem 19:2507–2511.

29. Ramis R, et al. (2019) A coarse-grained molecular dynamics approach to the study of the intrinsically disordered protein α-synuclein J Chem Inform Model 59:1458–1471.

30. Bhattacharya S, Xu L, Thompson D (2019) Molecular simulations reveal terminal group mediated stabilization of helical conformers in both amyloid-b42 and α-synuclein ACS Chem Neurosci 10:2830–2842.

31. Ilie IM, Caflisch A (2019) Simulation studies of amyloidogenic polypeptides and their aggregates Chem Rev 119:6956–6993.

32. Pietrek LM, Stelzl LS, Hummer G (2020) Hierarchical ensembles of intrinsically disordered proteins at atomic resolution in molecular dynamics simulations J Chem Theor Comput 16:725–737.

33. Braun AR, et al. (2012) α-Synuclein induces both positive mean curvature and negative gaussian curvature in membranes J Amer Chem Soc 134:2613–2620.

34. Braun AR, et al. (2014) α-Synuclein-induced membrane remodeling is driven by binding affinity, partition depth, and interleaflet order asymmetry J Amer Chem Soc 136:9962–9972.

35. Braun AR, et al. (2017) α-synuclein’s uniquely long amphipathic helix enhances its membrane binding and remodeling capacity J Membr Biol 250:183–193.

36. Nepal B, Leveritt J, Lazaridis T (2018) Membrane curvature sensing by amphipathic helices: Insights from implicit membrane modeling Biophys J 114:2128–2141.

37. Liu YL, et al. (2018) Molecular simulation aspects of amyloid peptides at membrane interface Biochim Biophys Acta-Biomembranes 1860:1906–1916.

38. Press-Sandler O, Miller Y (2018) Molecular mechanisms of membrane-associated amyloid aggregation: Computational perspective and challenges Biochim Biophys Acta-Biomembranes 1860:1889–1905.

39. Sahoo A, Matysiak S (2019) Computational insights into lipid assisted peptide misfolding and aggregation in neurodegeneration Phys Chem Chem Phys 21:22679–22694.

40. Hedger G, Sansom MSP (2016) Lipid interaction sites on channels, transporters and receptors: recent insights from molecular dynamics simulations Biochim Biophys Acta 1858:2390–2400.

41. Duncan AL, Song W, Sansom MSP (2020) Lipid-dependent regulation of ion channels and GPCRs: insights from structures and simulations Ann Rev Pharmacol Toxicol 60:31–50.

42. Lautenschlager J, et al. (2018) C-terminal calcium binding of α-synuclein modulates synaptic vesicle interaction Nature Comms 9.

43. Ouberai MM, et al. (2013) α-synuclein senses lipid packing defects and induces lateral expansion of lipids leading to membrane remodeling J Biol Chem 288:20883–20895.

44. Stansfeld PJ, Sansom MSP (2011) From coarse-grained to atomistic: a serial multi-scale approach to membrane protein simulations J Chem Theor Comp 7:1157–1166.

45. Fusco G, et al. (2014) Direct observation of the three regions in α-synuclein that determine its membrane-bound behaviour Nature Comms 5:3827.

46. Runfola M, et al. (2020) The N-terminal Acetylation of α-Synuclein Changes the Affinity for Lipid Membranes but not the Structural Properties of the Bound State Scientific Reports 10:204.

47. Harrigan MP, et al. (2017) MSMBuilder: Statistical models for biomolecular dynamics Biophys J 112:10–15.

48. Bowman GR, Huang XH, Pande VS (2009) Using generalized ensemble simulations and Markov state models to identify conformational states Methods 49:197–201.

49. Li YW, et al. (2018) Amyloid fibril structure of α-synuclein determined by cryoelectron microscopy Cell Res 28:897–903.

50. Li BS, et al. (2018) Cryo-EM of full-length α-synuclein reveals fibril polymorphs with a common structural kernel Nature Comms 9.

51. Pujols J, et al. (2018) Small molecule inhibits α-synuclein aggregation, disrupts amyloid fibrils, and prevents degeneration of dopaminergic neurons Proc Natl Acad Sci USA 115:10481–10486.

52. Wassenaar TA, et al. (2015) Computational lipidomics with *insane*: a versatile tool for generating custom membranes for molecular simulations J Chem Theor Comput 11:2144–2155.

53. Abraham MJ, et al. (2015) GROMACS: High performance molecular simulations through multi-level parallelism from laptops to supercomputers SoftwareX 1–2:19–25.

54. Hess B, Kutzner C, van der Spoel D, Lindahl E (2008) GROMACS 4: algorithms for highly efficient, load-balanced, and scalable molecular simulation J Chem Theor Comp 4:435–447.

55. Monticelli L, et al. (2008) The MARTINI coarse grained force field: extension to proteins J Chem Theor Comp 4:819–834.

56. Berendsen HJC, et al. (1984) Molecular dynamics with coupling to an external bath J Chem Phys 81:3684–3690.

57. Hess B, Bekker H, Berendsen HJC, Fraaije JGEM (1997) LINCS: A linear constraint solver for molecular simulations J Comp Chem 18:1463–1472.

58. Huang J, MacKerell AD (2013) CHARMM36 all-atom additive protein force field: Validation based on comparison to NMR data J Comp Chem 34:2135–2145.

59. Darden T, York D, Pedersen L (1993) Particle mesh Ewald - an N.log(N) method for Ewald sums in large systems J Chem Phys 98:10089–10092.

60. Bussi G, Donadio D, Parrinello M (2007) Canonical sampling through velocity rescaling J Chem Phys 126:014101.

61. Parrinello M, Rahman A (1981) Polymorphic transitions in single-crystals - a new molecular-dynamics method J Appl Phys 52:7182–7190.

62. Stansfeld PJ, Sansom MSP (2011) From coarse grained to atomistic: a serial multiscale approach to membrane protein simulations J Chem Theor Comput 7:1157–1166.

63. Humphrey W, Dalke A, Schulten K (1996) VMD-Visual Molecular Dynamics J Molec Graph 14:33–38.

64. Hunter JD (2007) Matplotlib: A 2D graphics environment Comput Sci Eng 9:90–95.

65. Pedregosa F, et al. (2011) Scikit-learn: Machine Learning in Python J Machine Learning Res 12:2825–2830.

66. Wrasidlo W, et al. (2016) A de novo compound targeting α-synuclein improves deficits in models of Parkinson’s disease Brain 139:3217–3236.

67. Marsh JA, Singh VK, Jia ZC, Forman-Kay JD (2006) Sensitivity of secondary structure propensities to sequence differences between α- and γ-synuclein: Implications for fibrillation Protein Sci 15:2795–2804.

68. Pervushin K, Riek R, Wider G, Wuthrich K (1997) Attenuated T2 relaxation by mutual cancellation of dipole-dipole coupling and chemical shift anisotropy indicates an avenue to NMR structures of very large biological macromolecules in solution Proc Natl Acad Sci USA 94:12366–12371.

69. Rappsilber J, Mann M, Ishihama Y (2007) Protocol for micro-purification, enrichment, pre-fractionation and storage of peptides for proteomics using StageTips Nature Protocols 2:1896–1906.

70. Tyanova S, Temu T, Cox J (2016) The MaxQuant computational platform for mass spectrometry-based shotgun proteomics Nature Protocols 11:2301–2319.

71. Yang B, et al. (2012) Identification of cross-linked peptides from complex samples Nature Methods 9:904–906.

72. Yuan ZF, et al. (2012) pParse: A method for accurate determination of monoisotopic peaks in high-resolution mass spectra Proteomics 12:226–235.

73. Wickham H (2009) ggplot2: Elegant Graphics for Data Analysis.

